# Statins attenuate antiviral IFN-β and ISG expression via inhibiting IRF3/JAK/STAT signaling in poly(I:C)-treated hyperlipidemic mice and macrophages

**DOI:** 10.1101/2020.06.21.163873

**Authors:** Atsushi Koike, Kaito Tsujinaka, Ko Fujimori

**Affiliations:** Department of Pathobiochemistry, Osaka University of Pharmaceutical Sciences, 4-20-1 Nasahara, Takatsuki, Osaka, 569-1094, Japan; Osaka University of Pharmaceutical Sciences, 4-20-1, Nasahara, Takatsuki, Osaka, 569-1094, Japan

**Author notes:** Corresponding author: Ko Fujimori.

**Keywords:** macrophage, Toll-like receptor (TLR), interferon, interferon regulatory factor (IRF), Janus kinase (JAK), signal transducers and activators of transcription 1 (STAT1), statin, interferon stimulated gene

## Abstract

Viral infection is a significant burden to healthcare worldwide. Statins, 3-hydroxy-3-methyl glutaryl coenzyme A reductase inhibitors, are widely used as cholesterol-lowering drugs. Recently, long term statin therapy was shown to reduce the antiviral immune response; however, the underlying molecular mechanisms are unclear. Here, we found that simvastatin decreased polyinosinic-polycytidylic acid [poly(I:C)]-induced expression of antiviral interferon (IFN)-β and IFN-stimulated genes (ISGs) in the bronchoalveolar lavage fluid and lungs of mice with high-fat diet-induced hyperlipidemia. As macrophages were the dominant cell type in the bronchoalveolar lavage fluid of poly(I:C)-treated mice, we examined the molecular mechanisms of statin-mediated inhibition of antiviral gene expression using murine J774.1/JA-4 macrophages. Simvastatin and pitavastatin decreased poly(I:C)-induced expression of IFN-β and ISGs. Moreover, they repressed poly(I:C)-induced phosphorylation of IFN regulatory factor (IRF) 3 and signal transducers and activators of transcription (STAT) 1, which is involved in Janus kinase (JAK)/STAT signaling. Mevalonate and geranylgeranylpyrophosphate (GGPP), but not cholesterol, counteracted the negative effect of statins on IFN-β and ISG expression and phosphorylation of IRF3 and STAT1. These results suggest that statins suppressed the expression of IFN-β and ISGs in poly(I:C)-treated hyperlipidemic mice and murine macrophages, and that these effects occured through the inhibition of IRF3-mediated JAK/STAT signaling in macrophages. Furthermore, GGPP recovered the statin-suppressed IRF3/JAK/STAT signaling pathway in poly(I:C)-treated macrophages.

## Introduction

Statins were discovered as a by-product in the research for new antimicrobial agents (1). They are competitive inhibitors of 3-hydroxy-3-methylglutaryl coenzyme A (HMG-CoA) reductase, the rate-limiting enzyme in the mevalonate pathway for the biosynthesis of cholesterol and isoprenoids (2). Statins are cholesterol-lowering drugs, which are widely used to treat dyslipidemia and reduce the risk of atherosclerotic cardiovascular disease (3). Recent studies indicate that statins also display lipid-independent “pleiotropic” effects that involve inhibiting the thrombogenic response, enhancing the stability of the atherosclerotic plaque, and decreasing the oxidative stress (4). Furthermore, the modification of inflammatory effects by statins has been demonstrated. Atorvastatin reduced the production of proinflammatory cytokines, e.g., tumor necrosis factor (TNF)-α and interleukin (IL)-6 via inhibiting nuclear factor-κB (NF-κB) signaling in vascular smooth muscle cells and mononuclear cells (5). In human hepatocytes, pravastatin, simvastatin, and atorvastatin exerted anti-inflammatory effects through the Janus kinase (JAK)/signal transducers and activators of transcription (STAT) signaling pathway (6). In human macrophages, the anti-inflammatory action of simvastatin was mediated by enhancing the expression of Kruppel-like factor 2 (7). Moreover, simvastatin suppressed the inflammatory response induced by polyinosinic-polycytidylic acid [poly(I:C)], a synthetic double-stranded RNA (dsRNA) analog, in bronchial epithelial cells (8). Thus, statins exert broad anti-inflammatory and immunomodulatory effects, possibly through cell type-specific mechanisms. At present, the mechanisms regulating statin-mediated modulation of the antiviral immune response in macrophages are not fully understood.

Viral infection is a significant burden to human and animal health. The innate immune cells, such as macrophages and dendritic cells, are the first line of defense against pathogens (9). In the case of viral infection, they detect a variety of viral components, including DNA, single- and doublestranded RNA, and glycoproteins, via pattern recognition receptors, such as Toll-like receptors (TLRs) and retinoic acid-inducible gene (RIG)-I-like receptors (RLRs) (10). dsRNA generated in the process of viral replication is recognized by TLR3 and RLRs, such as RIG-I and melanoma differentiation-associated gene 5 (MDA5), which activates the NF-κB and IFN regulatory factor (IRF) 3 signaling pathways and, in turn, enhances the production of pro-inflammatory cytokines, type I interferons (IFN-α and IFN-β) and a variety of IFN-stimulated genes (ISGs), such as CCL5/RANTES and viperin, to defend against viral infection (11–13). Moreover, the binding of IFNs to cognate receptors activates JAK/STAT signaling to induce the expression of various ISGs (14,15). Deficiency of IFNs, IFN receptors, or JAK/STAT signaling pathway enhances viral replication (16–18). Thus, IFNs and ISGs are critical regulators of the antiviral immune response.

In this study, we examined the effect of simvastatin and pitavastatin on poly(I:C)-induced expression of antiviral IFN-β and ISGs in murine macrophages and mice with high-fat diet (HFD)-induced hyperlipidemia, and investigated the underlying molecular mechanisms.

## Results

### Simvastatin attenuated poly(I:C)-induced IFN-β and ISG expression in mice with HFD-induced hyperlipidemia

HFD induces hyperlipidemia and hyperlipidemic mice have altered immune responses (23,24). Thus, we analyzed the effect of statins on the antiviral immune response in mice with HFD-induced hyperlipidemia. To induce hyperlipidemia, C57BL/6J mice were fed an LFD or HFD for 8 weeks. Then, the HFD-fed mice were administered simvastatin (30 mg/kg) once a day for the subsequent 10 days, followed by treatment with poly(I:C) for 8 h (Fig. 1a). Unlike LFD-fed mice, HFD-fed mice became obese (Fig. 1b). Serum levels of non-esterified fatty acid (NEFA), total cholesterol, low-density lipoprotein (LDL)-cholesterol, high-density lipoprotein (HDL)-cholesterol, and glucose, but not those of triglycerides, were higher in HFD-fed mice than in LFD-fed mice (Table 1). Simvastatin treatment reduced the levels of total- and LDL-cholesterol in HFD-fed mice (Table 1). These results indicate that the HFD caused a lipid imbalance in the serum and that simvastatin effectively lowered serum total- and LDL-cholesterol levels at 30 mg/kg.

**Figure 1.**
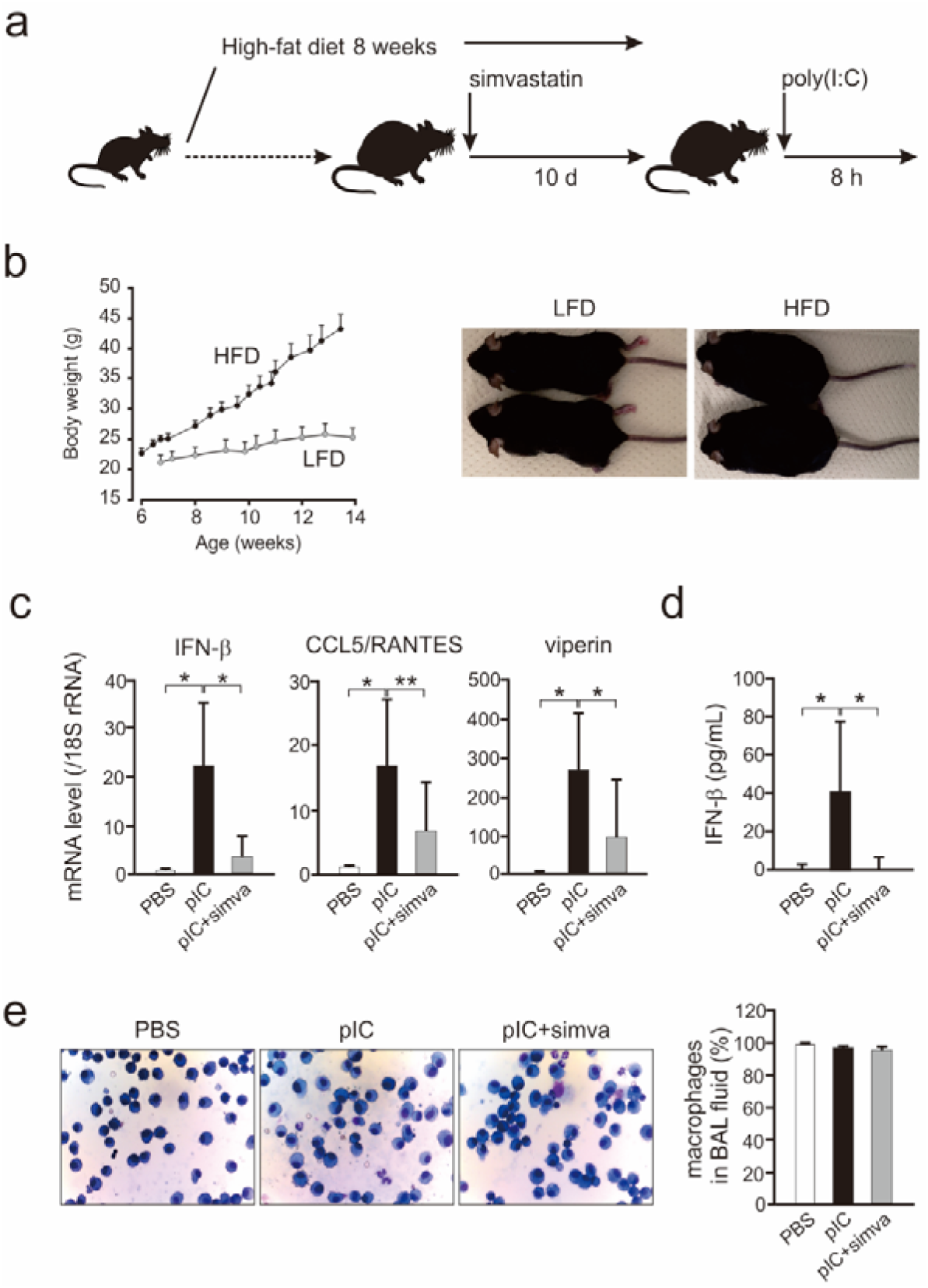
Simvastatin suppressed poly(I:C)-induced expression of IFN-β and ISGs in mice with HFD-induced hyperlipidemia. **a.** Schematic summary of the experimental design; **b.** Body weight changes in LFD-fed (gray circles; n=30) and HFD-fed (black circles; n=30) mice (left panel). Representative photographs of LFD- and HFD-fed mice (right panel); **c.** Changes to IFN-β and ISG expression in the lungs of mice treated with PBS (white columns), poly(I:C) (pIC; black columns), or poly(I:C) plus simvastatin (pIC+simva; gray columns) (n=10 in each group). Data are expressed as the fold change relative to PBS-treated mice, and shown as the mean ± SD, \**p* < 0.01, \*\**p* < 0.05; **d.** IFN-β level in the BAL fluid, as measured by ELISA (n=8-9 in each group). The results are presented as the mean ± SD, \**p* < 0.01, \*\**p* < 0.05; **e.** Left panel: Representative images of cells in the BAL fluid (Diff-Quik, x 400). Right panel: The percentage of macrophages to total cells in the BAL fluid of mice treated with PBS (white column), poly(I:C) (pIC; black column), or poly(I:C) plus simvastatin (pIC+simva; gray column) (n=4-5 in each group). The results are presented as the mean ± SD.

**Table 1.** Serum biochemical parameters in mice.

|  | After feeding<br>(8 weeks) |  | After treatment<br>(10 days) |  |
| --- | --- | --- | --- | --- |
|  | LFD<br>(n=30) | HFD<br>(n=30) | HFD+vehicle<br>(n=9) | HFD+simva<br>(n=10) |
| **NEFA (μEq/L)** | 1193.3±234.9 | 1697.2±248.9<sup>#</sup> | 1294.8±283.2 | 1296.6±218.0 |
| **Total cholesterol (mg/dL)** | 114.7±12.1 | 207.5±25.8<sup>#</sup> | 198.9±16.8 | 165.5±26.7<sup>*</sup> |
| **LDL-cholesterol (mg/dL)** | 11.8±2.0 | 18.6±6.1<sup>#</sup> | 18.2±3.3 | 10.6±3.6<sup>*</sup> |
| **HDL-cholesterol (mg/dL)** | 61.4±7.2 | 98.9±10.9<sup>#</sup> | 87.7±4.9 | 87.2±13.2 |
| **Glucose (mg/dL)** | 134.3±23.8 | 181.4±30.5<sup>#</sup> | 112.6±30.6 | 134.3±21.9 |
| **Triglycerides (mg/dL)** | 69.9±25.1 | 64.6±20.4 | 39.2±9.9 | 40.9±23.2 |

Values are expressed as the mean ± SD, <sup>#</sup>*p* < 0.01 vs. LFD-fed mice, <sup>\*</sup>*p* < 0.01 vs. vehicle-treated HFD-fed mice. LFD, low-fat diet; HFD, high-fat diet; NEFA, non-esterified fatty acids; LDL, low-density lipoprotein, HDL, high-density lipoprotein, simva, simvastatin.

We then examined the effects of simvastatin on the antiviral immune response in the lung. When HFD-fed mice were intranasally administered with poly(I:C), the mRNA levels of IFN-β, CCL5/RANTES, and viperin in the lungs were higher than those of PBS-treated mice (Fig. 1c). When simvastatin-treated HFD-fed mice were given poly(I:C), the mRNA levels of IFN-β, CCL5/RANTES, and viperin were approximately 85%, 61%, and 64% lower, respectively, than those in vehicle-treated HFD-fed mice (Fig. 1c). Moreover, the level of secreted IFN-β in the BAL fluid of mice with HFD-induced hyperlipidemia was increased after poly(I:C) exposure; this effect was clearly attenuated by simvastatin treatment (Fig. 1d). In addition, the majority of the cells in the BAL fluid of PBS-, poly(I:C)-, and poly(I:C) plus simvastatin-treated mice comprised macrophages (over 95% of total cells) (Fig. 1e). These results suggest that simvastatin decreased the expression of antiviral IFN-β and ISGs in the lungs of mice with HFD-induced hyperlipidemia.

### Effects of statins on cell viability and intracellular cholesterol level in macrophages

We investigated the molecular mechanisms underlying the above effects using cultured macrophages. First, we treated J774.1/JA-4 cells with different concentrations of simvastatin and pitavastatin for 48 h to evaluate their cytotoxicity (Fig. S1). Neither statin was cytotoxic at up to 100 nM, and both showed cytotoxicity at over 1 μM (Fig. 2a).

**Figure 2.**
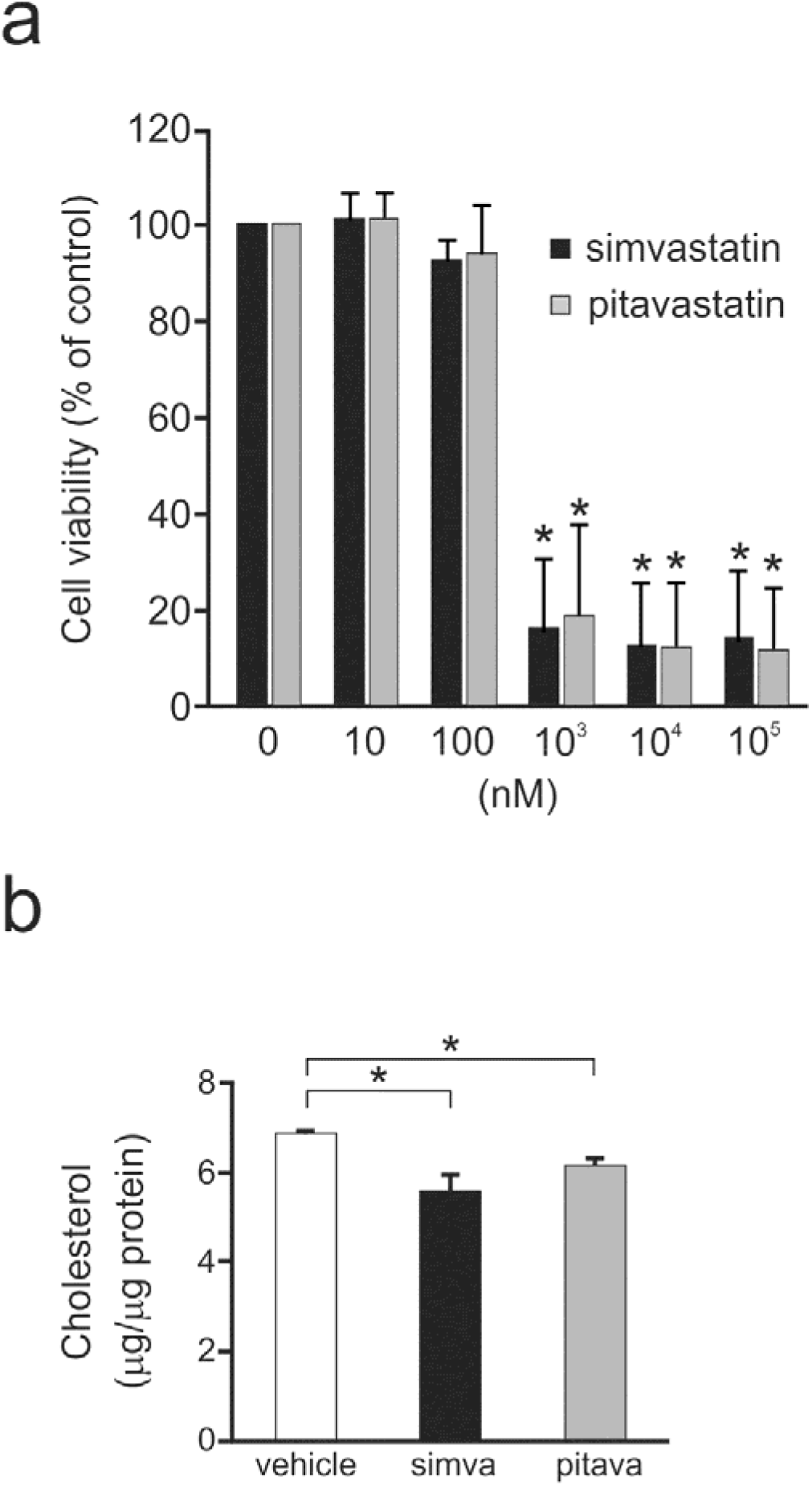
The effects of statins on cell viability and intracellular cholesterol level in macrophages. **a.** Cell viability assay. J774.1/JA-4 cells were treated with 0–100 μM of simvastatin (black columns) or pitavastatin (gray columns) for 48 h. Data are expressed as the fold change relative to untreated cells (0 nM), and shown as the mean ± SD from three independent experiments, \**p* < 0.01, vs. untreated cells (0 nM); **b.** Quantification of intracellular cholesterol levels in J774.1/JA-4 cells treated with vehicle (white column), simvastatin (simva; black column), or pitavastatin (pitava; gray column) (100 nM each) for 48 h. Intracellular cholesterol levels are expressed as μg of total cholesterol per μg of protein. The results are presented as the mean ± SD from three independent experiments, \**p* < 0.01.

Subsequently, we tested whether statins affect intracellular cholesterol levels in J774.1/JA-4 cells. Exposure to simvastatin or pitavastatin for 48 h decreased intracellular cholesterol levels by approximately 19% and 11%, respectively (Fig. 2b). Taken together, these results imply that 100 nM simvastatin or pitavastatin reduced intracellular cholesterol levels in macrophages without any cytotoxic effects. Thus, this dose was used in subsequent experiments.

### Statins attenuated poly(I:C)-induced expression of IFN-β and ISGs in macrophages

We examined the time-course of the effect of statins on the expression of IFN-β, ISGs (CCL5/RANTES, and viperin), and inflammatory cytokines (IL-6 and TNF-α) in macrophages. To this end, J774.1/JA-4 cells were treated with simvastatin or pitavastatin for 1, 20, or 48 h followed by treatment with poly(I:C). Expression of the antiviral cytokine genes was unchanged or weakly suppressed at the 1-h and 20-h mark, and markedly inhibited at the 48-h time point (Fig. 3a, Fig. S2). Thus, we focused on the effects of statins after 48 h of treatment.

**Figure 3.**
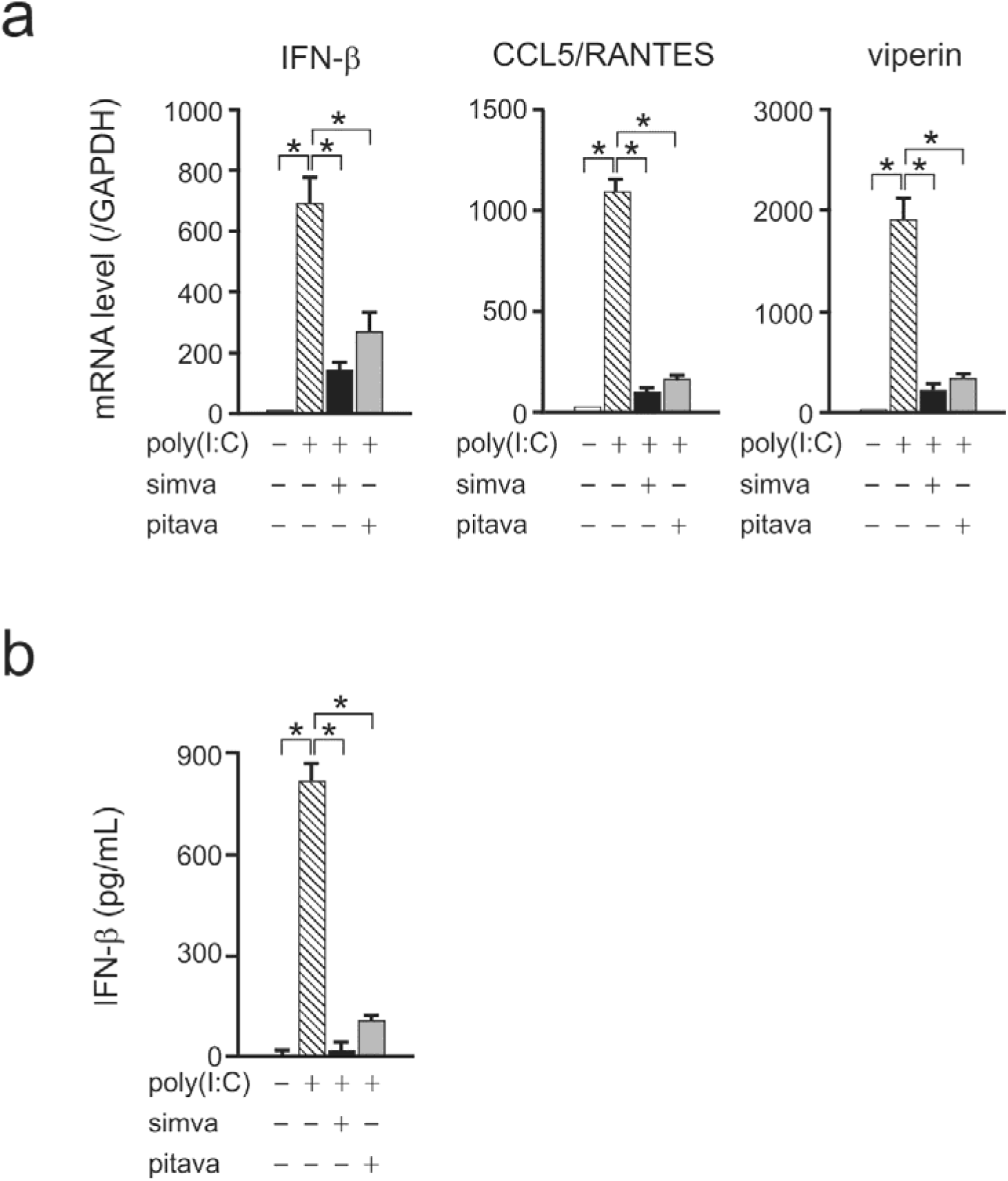
Statins attenuated the expression of IFN-β and ISGs in poly(I:C)-treated macrophages. **a.** Changes to gene expression in J774.1/JA-4 cells after 48 h incubation with vehicle (hatched columns), simvastatin (simva; black columns), or pitavastatin (pitava; gray columns) (100 nM each) followed by treatment with 10 μg/mL of poly(I:C) for 4 h. Data are expressed as the fold change relative to untreated cells (white columns). The results are presented as the mean ± SD from at least three independent experiments, \**p* < 0.01; **b.** Quantification of secreted IFN-β protein. J774.1/JA-4 cells (white column) were incubated with vehicle (hatched column), simvastatin (simva; black column), or pitavastatin (pitava; gray column) (100 nM each) for 48 h before they were treated with 10 μg/mL of poly(I:C) for 20 h. The results are presented as the mean ± SD from three independent experiments, \**p* < 0.01.

The mRNA levels of IFN-β, CCL5/RANTES, and viperin were increased by poly(I:C) treatment (Fig. 3a). Following a 48-h incubation in statin-containing medium, the mRNA levels of IFN-β, CCL5/RANTES, and viperin were decreased by 79%, 91%, and 86%, respectively, in simvastatin-exposed cells, and by 61%, 85%, and 83%, respectively, in pitavastatin-exposed cells (Fig. 3a). Moreover, each statin decreased poly(I:C)-induced IFN-β production by more than 90% (Fig. 3b). These results suggest that simvastatin and pitavastatin impaired poly(I:C)-induced expression of antiviral IFN-β and ISGs, and production of IFN-β in macrophages.

The mRNA levels of IL-6 and TNF-α were increased by poly(I:C) treatment (Fig. 4a). After a 48-h incubation in statin-containing medium, the mRNA levels of IL-6 and TNF-α were decreased by 65% and 36%, respectively, in simvastatin-treated cells, and by 49% and 26%, respectively, in pitavastatin-treated cells (Fig. 4a). Additionally, simvastatin reduced the production of IL-6 and TNF-α by 28% and 39%, respectively (Fig. 4b). Pitavastatin lowered TNF-α production by 22% without affecting IL-6 levels (Fig. 4b). These results reveal that statins impaired the gene and protein expression of inflammatory cytokines in poly(I:C)-treated macrophages.

**Figure 4.**
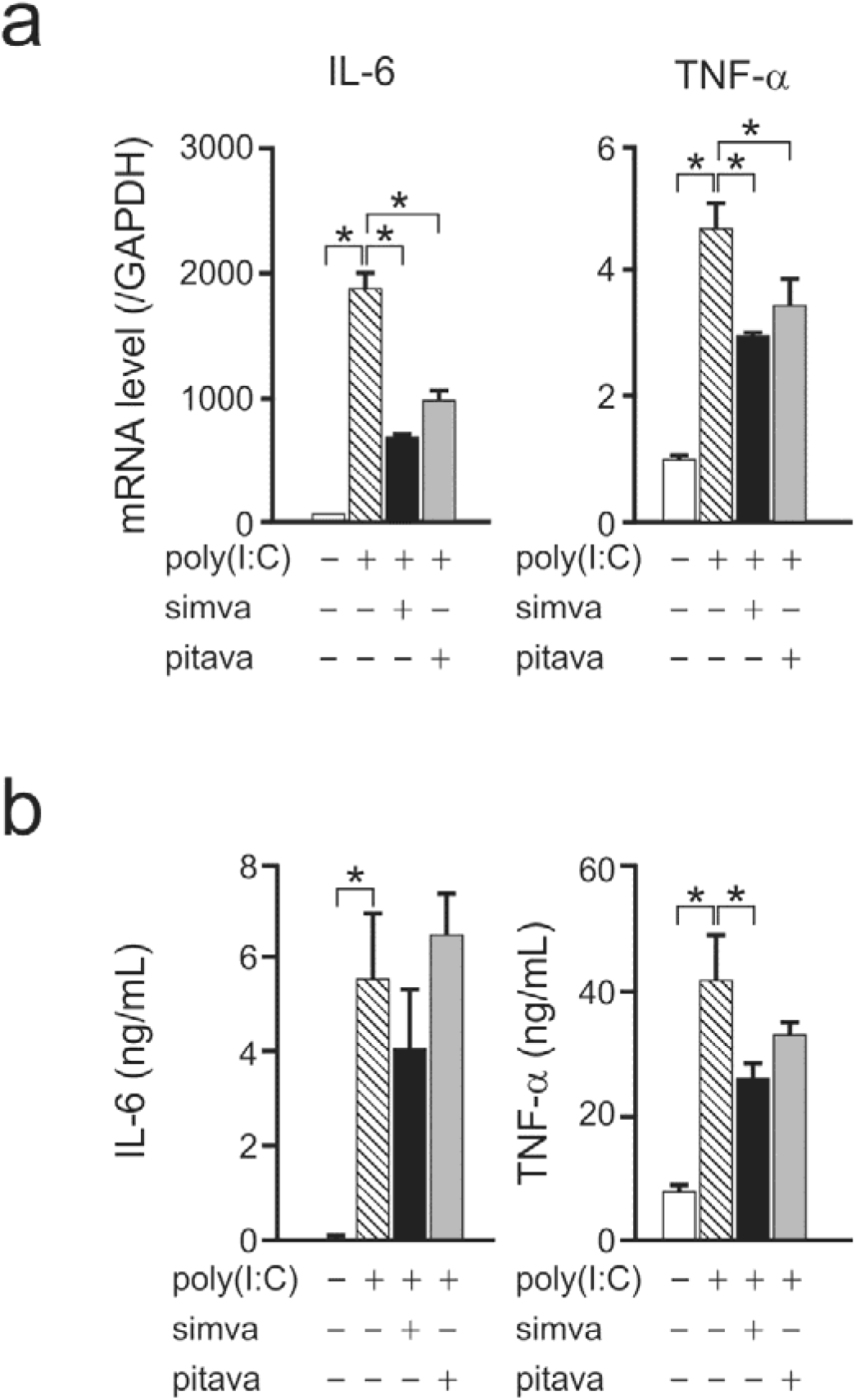
Statins impaired the expression of inflammatory cytokines in poly(I:C)-treated macrophages. **a.** Changes to the expression of inflammatory cytokine genes. J774.1/JA-4 cells were cultured and treated as described in the legend of Fig. 3a. Data are shown as the fold change relative to vehicle-treated cells. The results are presented as the mean ± SD from at least three independent experiments, \**p* < 0.01; **b.** Quantification of inflammatory cytokine levels. J774.1/JA-4 cells were cultured and treated as described in the legend of Fig. 3b. The results are presented as the mean ± SD from three independent experiments, \**p* < 0.01.

### Statins did not affect the cellular uptake of poly(I:C) by macrophages

Double-stranded RNA and poly(I:C) are recognized by intracellular receptors, such as TLR3, RIG-I, and MDA5 (10); thus, we examined whether statins could affect the cellular uptake of poly(I:C). J774.1/JA-4 cells were cultured with simvastatin or pitavastatin for 48 h, and then further incubated with poly(I:C) (HMW)-Rhodamine for 2 h. The uptake of rhodamine-labelled poly(I:C) into cells did not differ between vehicle- and statin-treated cells (Fig. 5). In addition, the intracellular poly(I:C) (HMW)-Rhodamine/Hoechst fluorescent ratio was similar between vehicle- and statin-treated cells (Fig. 5). These observations indicate that statins did not affect the uptake of poly(I:C) by macrophages.

**Figure 5.**
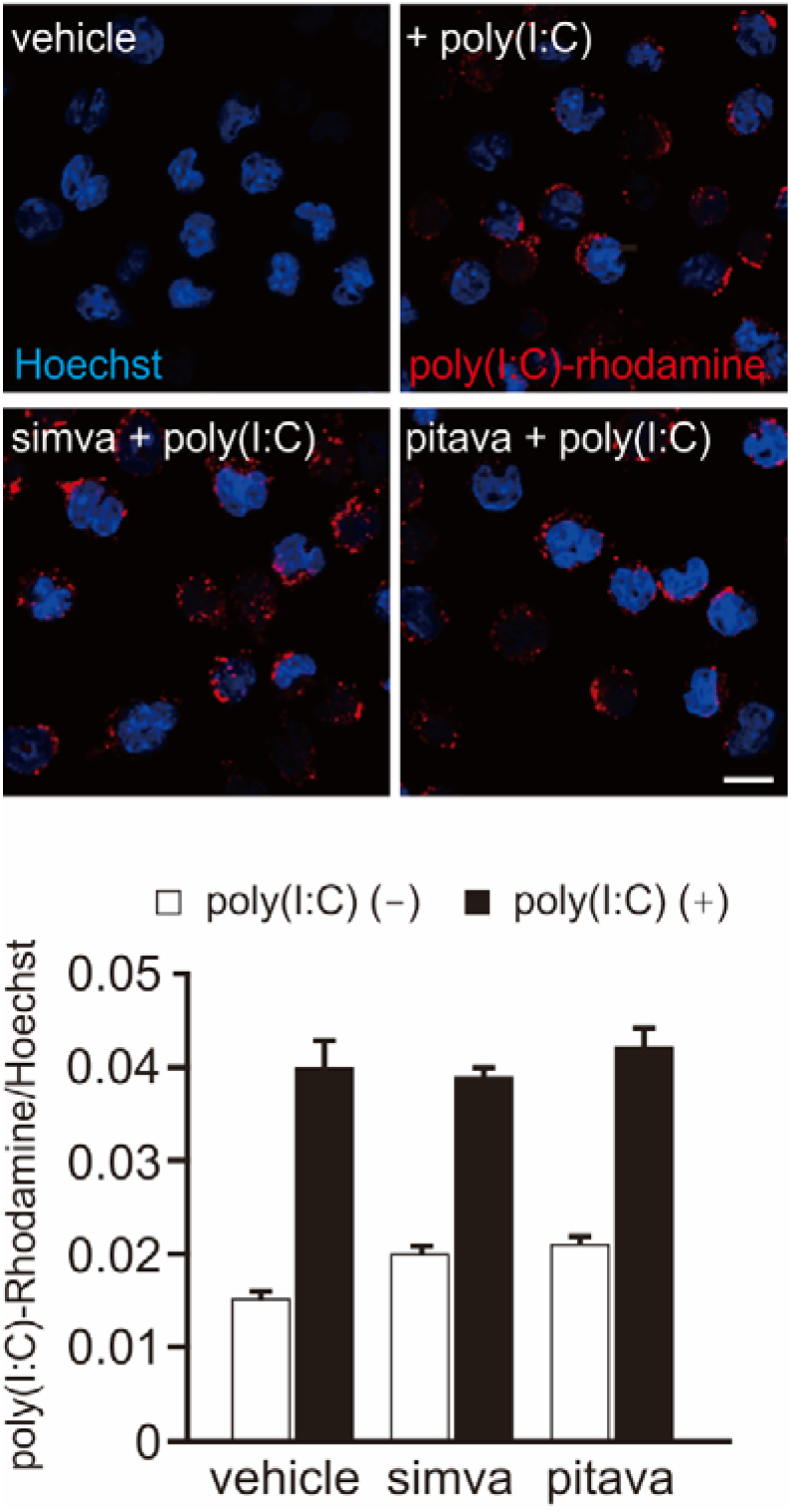
Statins did not affect the cellular uptake of poly(I:C). J774.1/JA-4 cells were cultured with simvastatin (simva) or pitavastatin (pitava) (100 nM each) for 48 h, then treated with (black columns) or without (white columns) 1 μg/mL of rhodamine-labeled poly(I:C) (red) for 2 h. The cells were also stained with Hoechst 33342 (blue). Fluorescence intensity of rhodamine-labeled poly(I:C) was normalized to that of Hoechst 33342. Scale bar = 10 μm. The results are presented as the mean ± SD from three independent experiments.

### Poly(I:C) enhanced IFN-β and ISG expression through TLR3 in macrophages

A previous study reported that the expression of RIG-I and MDA5 is induced by TLR3 signaling, suggesting that TLR3 is the first receptor to sense dsRNA and poly(I:C) (25). TLR3 expression levels were comparable between vehicle-, simvastatin-, and pitavastatin-treated macrophages (Fig. 6a). To verify the involvement of TLR3 in the regulation of antiviral IFN-β and ISGs expression in poly(I:C)-treated macrophages, we knocked down TLR3 expression using siRNA. TLR3 siRNA decreased TLR3 mRNA levels to 74% of that of negative control siRNA (Fig. 6b). siRNA-mediated TLR3 knockdown reduced the transcription of IFN-β, CCL5/RANTES, and viperin genes (Fig. 6c). These results imply that TLR3 is an upstream regulator of the expression of antiviral IFN-β and ISGs in poly(I:C)-treated macrophages.

**Figure 6.**
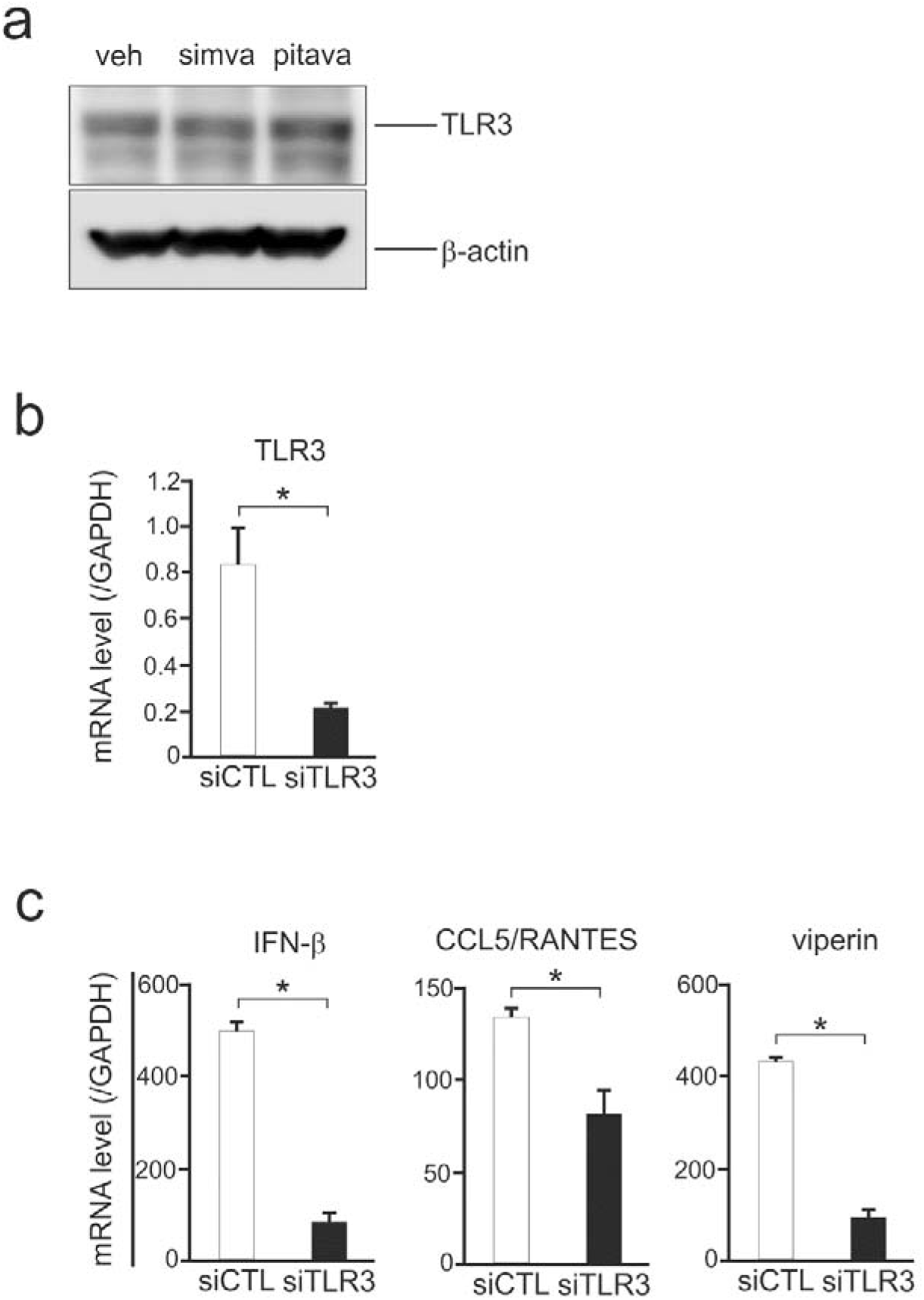
The effects of statins on the expression of TLR3 and involvement of TLR3 in the expression of IFN-β and ISGs in poly(I:C)-treated macrophages. **a.** Western blot analysis for TLR3. J774.1/JA-4 cells were cultured with simvastatin or pitavastatin (100 nM) for 48 h then subjected to western blot analysis using 25 μg of protein in each lane. β-actin was used as the internal control. The results are representative of three independent experiments; **b.** Efficiency of TLR3 knockdown. J774.1/JA-4 cells were transfected with TLR3 siRNA (siTLR3; 20 nM, black column) or negative control siRNA (siCTL; 20 nM, white column), and incubated for 24 h. The results are presented as the mean ± SD from at least three independent experiments, \**p* < 0.01; **c.** Changes to the expression of IFN-β and ISGs after siRNA treatment. siTLR3-transfected (black columns) and negative control siRNA-transfected (white columns) J774.1/JA-4 cells were treated with 10 μg/mL of poly(I:C) for 4 h. The results are presented as the mean ± SD from at least three independent experiments, \**p* < 0.01.

### Statins inhibited IRF3-mediated JAK/STAT signaling in poly(I:C)-treated macrophages

TLR3 activates IRF3, which promotes the expression of antiviral IFN-β and ISGs and activates the JAK/STAT signaling in macrophages (26). Thus, we examined whether statins affect IRF3-mediated JAK/STAT signaling in poly(I:C)-exposed macrophages. In vehicle-treated cells, phosphorylation (activation) of IRF3 was enhanced 30 and 60 min after exposure to poly(I:C), before gradually decreasing until the 180-min mark; phosphorylation of STAT1 was increased 180 min after exposure to poly(I:C) (Fig. 7a). Simvastatin and pitavastatin inhibited the phosphorylation of both IRF3 and STAT1 in poly(I:C)-treated macrophages (Fig. 7a). Inhibition of JAK, the upstream activator of STAT, with tofacitinib impaired poly(I:C)-induced phosphorylation of STAT1 (Fig. 7b) and decreased the gene expression of CCL5/RANTES and viperin by 11% and 25%, respectively. The expression level of IFN-β was lowered only slightly by tofacitinib (Fig. 7c). These findings suggest that statins inhibited IRF3-mediated JAK/STAT signaling in poly(I:C)-exposed macrophages.

**Figure 7.**
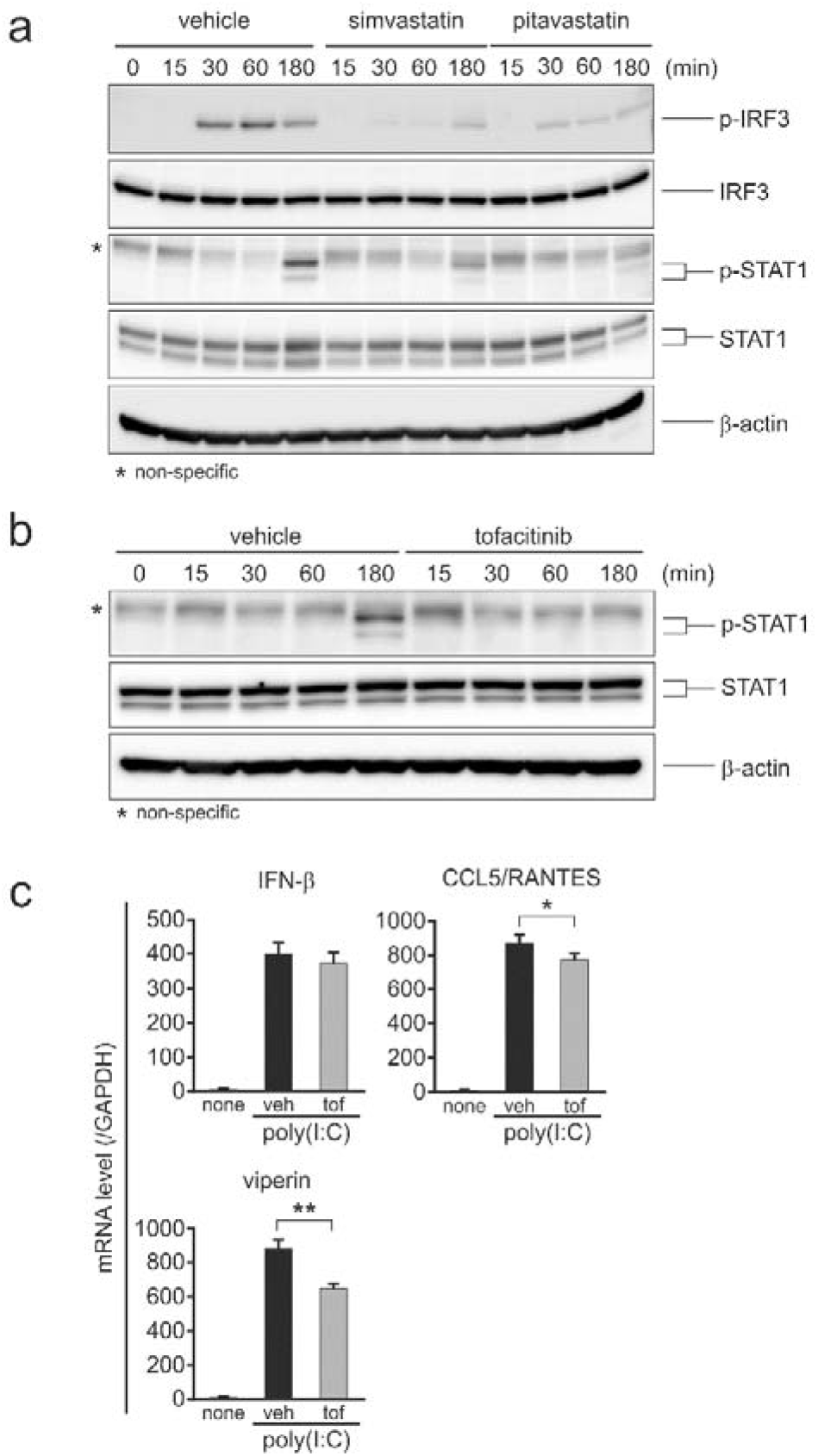
Statins inhibited IRF3-mediated JAK/STAT signaling in poly(I:C)-treated macrophages. **a.** Western blot analysis for IRF3 and STAT1. J774.1/JA-4 cells were cultured with simvastatin or pitavastatin (100 nM each) for 48 h, then treated with 10 μg/mL of poly(I:C) for the indicated time periods. An equal amount of protein (15 μg) was applied in each lane. The results are representative of three independent experiments. β-actin was used as the internal control; **b.** The effect of tofacitinib (JAK inhibitor) on poly(I:C)-induced activation of STAT1. J774.1/JA-4 cells were pre-treated with vehicle or 1 μM of tofacitinib for 1 h, then exposed to 10 μg/mL of poly(I:C) for the indicated time periods. The results are representative of three independent experiments, *non-specific signals that were detected in all samples; **c.** Changes to gene expression in tofacitinib-treated cells. J774.1/JA-4 cells were pre-treated with vehicle (veh; black columns) or 1 μM tofacitinib (tof; gray columns) for 1 h, then exposed to 10 μg/mL of poly(I:C) for 4 h. The mRNA levels are expressed as the fold change relative to untreated cells (white columns). The results are presented as the mean ± SD of three independent experiments, \**p* < 0.01, \*\**p* < 0.05.

### GGPP counteracted the inhibitory effect of statins on IFN-β and ISG expression

Statins suppress the mevalonate pathway for the biosynthesis of cholesterol and isoprenoid intermediates, such as GGPP (12) (Fig. 8a). We investigated the role of the mevalonate pathway in the inhibitory effect of statins on IFN-β and ISG expression in J774.1/JA-4 cells. The cells were cultured with simvastatin or pitavastatin together with mevalonate, GGPP, or cholesterol, then treated with poly(I:C). Mevalonate and GGPP, but not cholesterol, prevented the statin-induced inhibition of IFN-β, CCL5/RANTES, and viperin gene expression, although neither mevalonate or GGPP alone affected the expression of these genes (Fig. 8b). Moreover, mevalonate and GGPP reversed the negative effect of statins on the production of IFN-β (Fig. 8c). These results suggest that GGPP rescued the statin-induced inhibition of antiviral IFN-β and ISG expression in poly(I:C)-treated macrophages.

**Figure 8.**
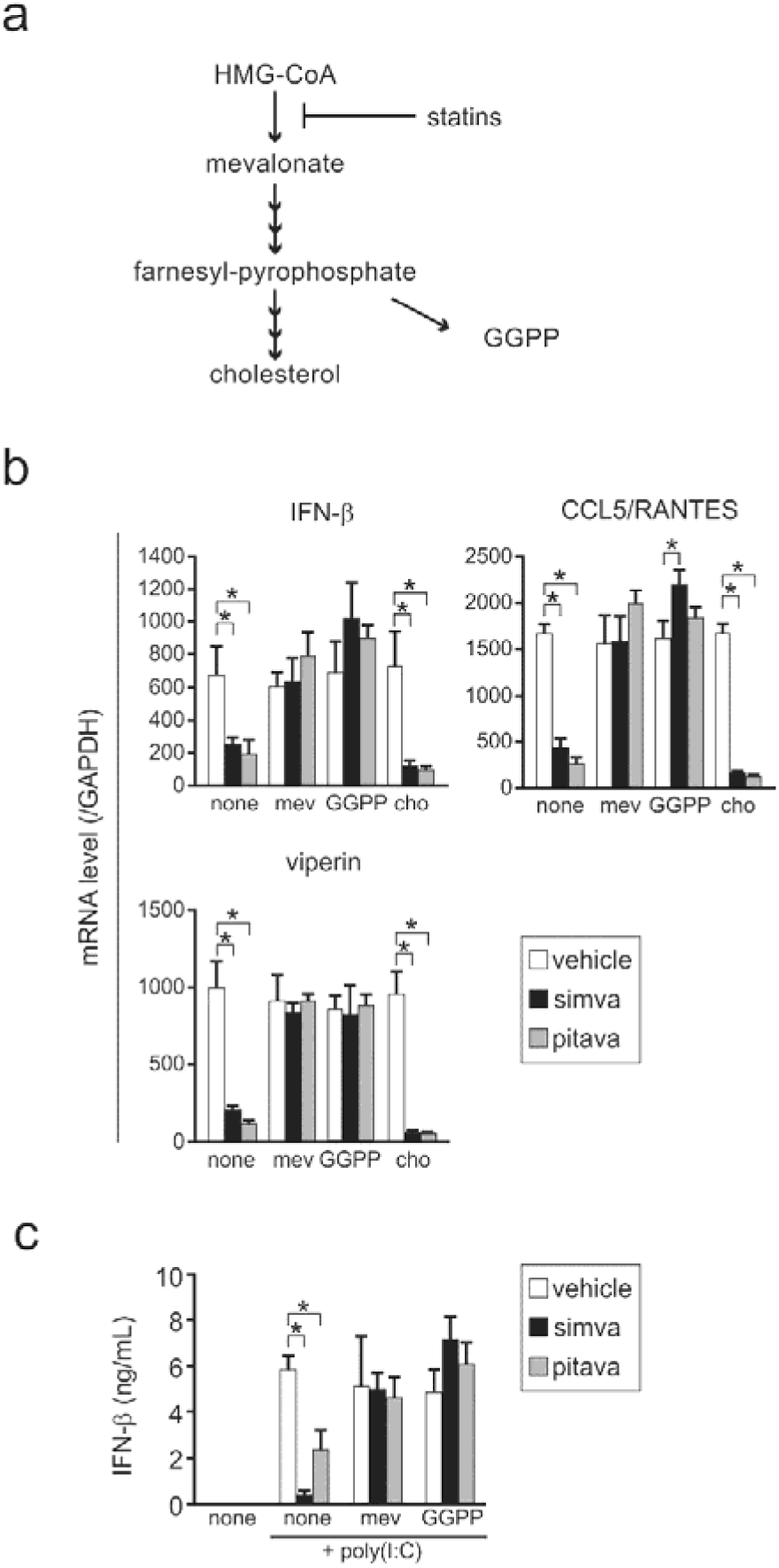
Mevalonate and GGPP reversed the inhibitory effect of statins on IFN-β and ISG expression in poly(I:C)-treated macrophages. **a.** Schematic representation of the cholesterol and isoprenoid biosynthesis pathway; **b.** Mevalonate and GGPP reversed the inhibitory effect of simvastatin and pitavastatin on IFN-β and ISG expression. J774.1/JA-4 cells (white columns) were cultured with simvastatin (simva; black columns) or pitavastatin (pitava; gray columns) for 24 h, then incubated with 300 μM of mevalonate (mev), 1 μg/mL of GGPP, or 1 μg/mL of cholesterol (cho) for 24 h. The cells were further treated with 10 μg/mL of poly(I:C) for 4 h. The mRNA levels are expressed as the fold change relative to untreated cells. The results are presented as the mean ± SD from three independent experiments, \**p* < 0.01; **c.** Quantification of IFN-β levels. The cells were treated as described in the legend of Fig. 8b, and incubated with 10 μg/mL of poly(I:C) for 20 h. The results are presented as the mean ± SD from three independent experiments, \**p* < 0.01.

### GGPP is involved in the activation of IRF3-mediated JAK/STAT signaling

Next, we examined the role of GGPP in the regulation of IRF3-mediated JAK/STAT signaling in poly(I:C)-treated macrophages. J774.1/JA-4 cells were cultured in the presence of simvastatin together with mevalonate or GGPP, then exposed to poly(I:C). The activation of IRF3 was enhanced 30 and 60 min after treatment with poly(I:C), before gradually decreasing until the 180-min mark; the activation of STAT1 was increased 180 min after treatment with poly(I:C) (Fig. 9). Poly(I:C)-mediated activation of IRF3 and STAT1 was suppressed by co-treatment with simvastatin, and this effect was counteracted by the addition of GGPP or mevalonate (Fig. 9). Thus, GGPP may be involved in the activation of IRF3-mediated JAK/STAT signaling in poly(I:C)-treated macrophages.

**Figure 9.**
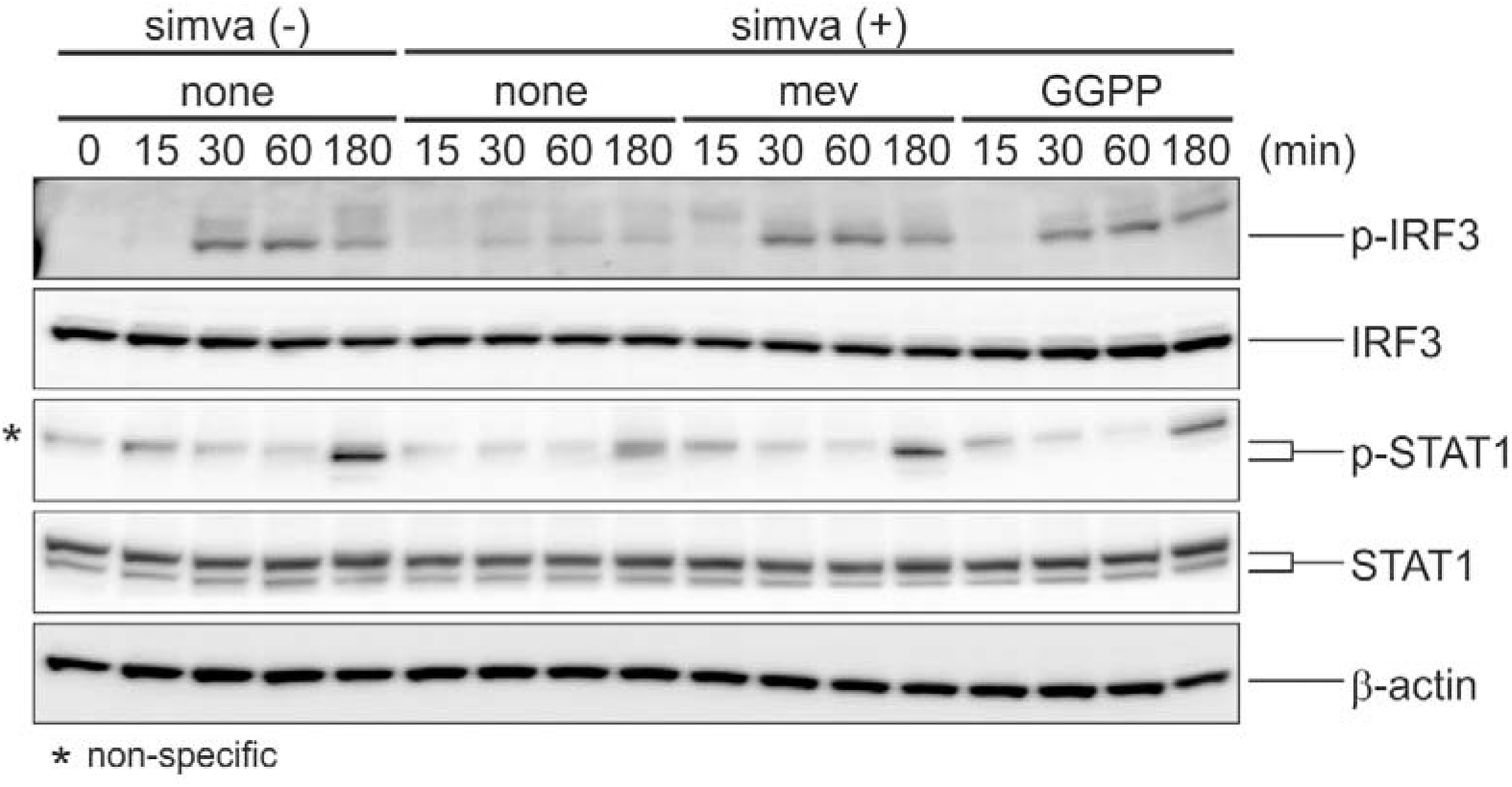
Mevalonate and GGPP reversed the inhibitory effect of simvastatin on IRF3-mediated JAK/STAT1 signaling in poly(I:C)-treated macrophages. Western blot analysis for IRF3 and STAT1. J774.1/JA-4 cells were cultured with simvastatin (simva), mevalonate (mev), and GGPP as described in the legend of Fig. 8b, then treated with 10 μg/mL of poly(I:C) for the indicated time periods. The results are representative of three independent experiments. β-actin was used as the internal control, *non-specific signals that were detected in all samples.

## Discussion

In this study, we found that statins suppressed the expression of antiviral IFN-β and ISGs in poly(I:C)-treated hyperlipidemic mice and murine macrophages, and that in macrophages these effects occurred via the inhibition of IRF3-mediated JAK/STAT signaling. GGPP counteracted the negative effect of statins on poly(I:C)-induced expression of IFN-β and ISGs and activation of IRF3 and STAT1 *in vitro*. Taken together, a schematic representation of how statins attenuate poly(I:C)-induced expression of antiviral IFN-β and ISGs is proposed in Fig. S3.

Statins, which are widely used to treat dyslipidemia and prevent cardiovascular diseases (3), are also known to exert anti-inflammatory and immunomodulatory actions (4). Long term statin therapy reduced the immune response to an influenza vaccine (27–29) and increased the risk of infection with herpes zoster virus (30,31). In airway epithelia, simvastatin suppressed poly(I:C)-induced inflammatory response through decreasing STAT3 activation and RANTES expression (8). The effects of statins on the antiviral immune response seem to be cell type-specific (5–8,32), but the mechanisms involved are not fully understood. Herein, we examined the effect of statins on the antiviral immune response in poly(I:C)-treated mice and macrophages, and investigated the underlying molecular mechanisms.

Hyperlipidemic mice exhibit altered immune responses (23,24,33). Atorvastatin improved plaque stability in ApoE-knockout mice (39), and exerted hypolipidemic and anti-inflammatory action in ApoE/LDL receptor-knockout mice (40). However, to our knowledge, the effects of statins on the antiviral immune response in hyperlipidemic mice have not been evaluated. Herein, we provided evidence that simvastatin decreases poly(I:C)-induced expression of antiviral IFN-β and ISGs in the lungs of mice with HFD-induced hyperlipidemia, and reduces IFN-β levels in their BAL fluid (Fig. 1c, d). Therefore, simvastatin modulates the antiviral response in hyperlipidemic mice. In a further study, we will employ a viral infection model rather than using a synthetic dsRNA mimic to confirm the effects of simvastatin *in vivo*.

Macrophages play a central role in innate immunity and serve as the first line of defense against pathogens, including bacteria and viruses (9). Depletion of macrophages in the lungs or brains of mice enhances their viral infectivity, leading to severe inflammation (34,35). Once macrophages have recognized the invading viruses, they express IFNs and ISGs to fight against the infection (12,13). IFNs also promote the proliferation of effector lymphocytes to provide long-lasting protection from specific viruses (36). Macrophages are the dominant cell type in the BAL fluid (37,38). In the present study, simvastatin and pitavastatin suppressed the expression of antiviral IFN-β and ISGs, decreased the production of IFN-β (Fig. 3a, b), and inhibited the gene and protein expression of inflammatory cytokines in poly(I:C)-treated J774.1/JA-4 cells (Fig. 4a, b). These findings suggest that statins impair the antiviral immune response in macrophages.

Activated IRF3 induces IFN-β gene expression which, in turn, activates JAK/STAT signaling, leading to the transcription of a variety of ISGs (39). Here, we showed that statins inhibit the phosphorylation of both IRF3 and STAT1 in poly(I:C)-treated macrophages (Fig. 7a, b), indicating that the negative effect of statins on the antiviral immune response occurs through the inhibition of IRF3-mediated JAK/STAT signaling in macrophages. In bronchial epithelial cells, simvastatin impairs dsRNA-induced IRF3 activation and IFN-β production, producing an antiinflammatory effect (40). Therefore, statins may inhibit the IRF3-mediated JAK/STAT signaling pathway in multiple cell types.

GGPP is a major isoprenoid intermediate in the mevalonate pathway. Blanco-Colio *et al*. showed that some of the pleiotropic effects of statins are suppressed by GGPP (41). In line with these results, we found that GGPP and mevalonate reversed the inhibitory effect of statins on the expression of antiviral IFN-β and ISGs and the phosphorylation of IRF3 and STAT1 (Figs. 8b, c and 9). GGPP is required for the geranylgeranylation of small G proteins, such as Rho, Rac, and Cdc42 (42). Inhibition of Rho and its downstream target, ROCK (Rho-associated, coiled-coil-containing protein kinase), is the principal mechanism behind the pleiotropic effects of statins (43). However, Y27632, a ROCK inhibitor, did not affect the expression of IFN-β or ISGs in poly(I:C)-treated macrophages (data not shown). Thus, the role of GGPP in statin-mediated suppression of the antiviral immune response in macrophages requires further investigation.

In conclusion, our data provides evidence that statins suppress the expression of antiviral IFN-β and ISGs in poly(I:C)-treated mice and macrophages, and that these effects are mediated through the inhibition of IRF3-mediated JAK/STAT signaling in macrophages. Moreover, we showed that GGPP counteracts the effects of statins on this signaling pathway in poly(I:C)-treated macrophages. Further studies using viral infection models are warranted to confirm this regulatory mechanism *in vitro* and *in vivo*.

## Experimental procedures

### Reagents

Simvastatin, pitavastatin, and DL-mevalonic acid lactone were purchased from Fujifilm Wako Pure Chemical (Osaka, Japan). The sodium salt of poly(I:C) and ammonium salt of geranylgeranyl pyrophosphate (GGPP) were obtained from Sigma Aldrich (St. Louis, MO, USA). Cholesterol and Hoechst 33342 were from Nacalai Tesque (Kyoto, Japan). Poly(I:C) (HMW)-Rhodamine was from InvivoGen (San Diego, CA, USA) and 4’,6-Diamidino-2-phenylindole dihydrochloride (DAPI) from Thermo Fisher Scientific (Waltham, MA, USA). The following antibodies were used in this study: anti-IRF3 (#4302), anti-phospho-IRF3 (#29047) (Cell Signaling Technology, Danvers, MA, USA), anti-TLR3 (AB62566; Abcam, Cambridge, UK), anti-phospho-STAT1 (p-STAT-1, Tyr701; AF2894; R&D Systems, Minneapolis, MN, USA) and anti-STAT1 (66545-1-Ig; Proteintech, Rosemont, IL, USA). Anti-β-actin antibody (A2228) was from Sigma Aldrich. Horseradish peroxidase (HRP)-linked anti-rabbit IgG (#7074) and anti-mouse IgG (sc-2371) were from Cell Signaling Technology and Santa Cruz Biotechnology (Dallas, TX, USA), respectively.

### Activation of simvastatin

Simvastatin (5 mg) was dissolved in 100 μL of ethanol before 150 μL of 0.1 M NaOH was added. The solution was incubated at 50 °C for 2 h and then neutralized to pH 7.0 with 1 M HCl. The resulting solution was diluted to 1 mL with distilled water, and aliquots were stored at −80 °C. The stock solution was diluted with phosphate-buffered saline (PBS) immediately before use.

### Animals and treatment

All animal experiments were performed in accordance with the ethical guidelines for the use of laboratory animals, and the study protocol was reviewed and approved by the Animal Committee of Osaka University of Pharmaceutical Sciences (No. 21). The animals were housed under controlled conditions (24±1 °C, humidity of 55±10%, and a 12 h light/dark cycle) and given *ad libitum* access to food and water. Male C57BL/6J mice (6 weeks old, n=60; Japan SLC; Shizuoka, Japan) were randomly divided into the following four groups: (1) low-fat diet (LFD) group (n=30; D12450B, Research Diet; New Brunswick, NJ, USA), (2) high-fat diet (HFD) (D12492, Research Diet) + PBS group (n=10), (3) HFD + poly(I:C) group (n=10), (4) HFD + simvastatin + poly(I:C) group (n=10). The HFD was a defined lard-based diet with 60% of its calories derived from fat, 20% from protein, and 20% from carbohydrates. The LFD was used as a defined control diet with identical nutritional composition as HFD, but different fat and carbohydrate content (10% of calories from fat, 20% from protein, and 70% from carbohydrates). The mice were fed the LFD or HFD for 8 weeks. Then, the HFD-fed mice were orally administered simvastatin (30 mg/kg) or vehicle (the same composition as the simvastatin solution, except for simvastatin) once a day for an additional 10 days. Statin dosage used in previous mouse and rat studies was 10–100 mg/kg per day (19–21). At 24 h after the final treatment, the mice were anesthetized by isoflurane inhalation, and 30 μL of poly(I:C) in PBS (1 mg/mL) was administered intranasally. At 8 h after administration, the bronchoalveolar lavage (BAL) fluid was collected under anesthesia. The BAL fluid was centrifuged at 400 x *g* for 3 min at 4 °C; the supernatant was used for the measurement of IFN-β levels while the cell pellet was resuspended in PBS and smeared using Cytospin 4 Cytocentrifuge at 400 x *g* for 5 min (Thermo Fisher Scientific). The cells were stained with Diff-Quik (Sysmex, Kobe, Japan) according to the manufacture’s protocols, then counted. The left lungs were removed for total RNA extraction as described below.

### Cell culture and treatment

Murine J774.1/JA-4 macrophages were cultured in Ham’s F-12 (Fujifilm Wako Pure Chemical) supplemented with 10% (v/v) heat-inactivated fetal bovine serum (FBS; Thermo Fisher Scientific) and 1% (v/v) antibiotic mixture (5000 U/mL penicillin and 5000 μg/mL streptomycin; Nacalai Tesque), and maintained at 37 °C in a humidified incubator with 5% CO_2_. J774.1/JA-4 cells were seeded in 60 mm dishes (2 x 10^6^ cells/dish) and treated with 100 nM of simvastatin or pitavastatin for 48 h. The cells were then harvested and seeded at 5 x 10^5^ cells/well in 12-well plates. Approximately 4 h after adhering to the plates, the cells were treated with 10 μg/mL of poly(I:C) for the indicated time periods.

### Evaluation of cell viability

Cell viability was assessed using the Cell Counting Kit-8 assay (CCK-8; Dojindo Molecular Technologies, Kumamoto, Japan) according to the manufacturer’s protocol. Briefly, J774.1/JA-4 cells were seeded at 5 x 10^4^ cells/well in 96-well plates, and then treated with various concentrations of simvastatin or pitavastatin for 0–48 h at 37 °C. After discarding the medium, 100 μL of fresh medium containing the CCK-8 solution (CCK-8: medium = 1:10; v/v) was added, and the cells were incubated for an additional 1 h at 37 °C. The absorbance of each well at 450 nm was measured using a Multiskan FC microplate photometer (Thermo Fisher Scientific).

### Western blot analysis

Cell lysates were prepared as described previously (22), and their protein content was measured using a BCA Protein Assay Kit (Thermo Fisher Scientific). The proteins were separated by SDS-PAGE, and then transferred onto PVDF membranes (Millipore, Bedford, MA, USA). The membranes were blocked with Immobilon Signal Enhancer (Millipore) for 1 h at 20 °C, followed by incubation with the primary antibodies in Immobilon Signal Enhancer for 1 h at 20 °C. After washing them with Tris-buffered saline containing 0.1% (v/v) Tween-20 (3 x 5 min), the membranes were probed with corresponding HRP-linked secondary antibodies in Immobilon Signal Enhancer for 1 h at 20 °C. Protein-antibody binding was visualized using Immobilon Forte Western HRP Substrate (Millipore) on the LAS-3000 imaging system (FUJIFILM, Tokyo, Japan).

### Quantitative PCR

Total RNA was extracted using ISOGEN II (Nippon Gene, Tokyo, Japan). First-strand cDNAs were synthesized using the ReverTra Ace^®^ qPCR RT Master Mix (Toyobo, Osaka, Japan) according to the manufacturer’s protocol. The cDNAs served as templates for quantitative PCR (qPCR) using Power SYBR^®^ Green Master Mix (Thermo Fisher Scientific) on the LightCycler 96 System (Roche Diagnostics, Mannheim, Germany). All genespecific primers are listed in Table 2. Target gene expression was calculated using the 2^−ΔΔCt^ method and normalized to that of glyceraldehyde-3-phosphate dehydrogenase (GAPDH) or 18S rRNA, which were used as internal controls.

**Table 2.**
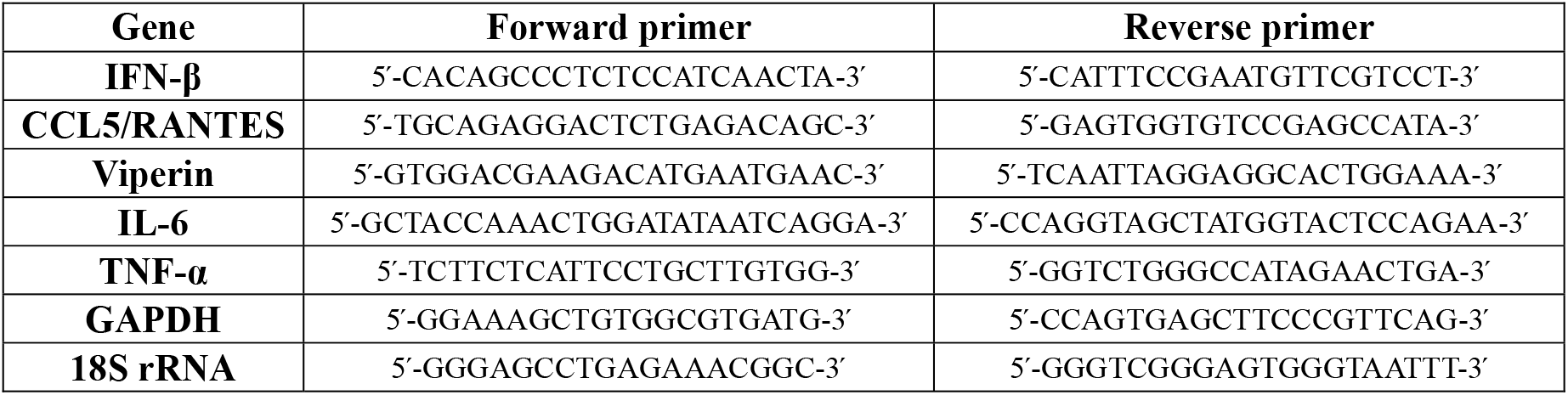
Nucleotide sequences of the primers used in qPCR.

### Measurement of IFN-β, IL-6, and TNF-α protein levels

J774.1/JA-4 cells were seeded at 5 x 10^5^ cells/well in 12-well plates. The levels of IFN-β, IL-6, and TNF-α in culture medium were determined using the Mouse IFN-β DuoSet ELISA, Mouse IL-6 Quantikine ELISA kit, and Mouse TNF-alpha Quantikine ELISA kit (R&D Systems), respectively, according to the manufacturer’s protocols.

### Quantification of intracellular cholesterol

J774.1/JA-4 cells (3 x 10^6^) were lysed with 0.3 mL of RIPA buffer [50 mM Tris-HCl, pH 8.0, 2 mM EDTA, pH 8.0, 150 mM NaCl, 0.5% (w/v) sodium deoxycholate, 0.1% (w/v) SDS, 1% (v/v) Nonidet P-40, 1% (v/v) Triton X-100, and 1% (v/v) protease inhibitor cocktail (Sigma)], followed by sonication (2 x 10 sec). Intracellular cholesterol levels were quantified with the Amplex Red Cholesterol Assay Kit (Thermo Fisher Scientific) according to the manufacturer’s instructions. Fluorescence intensity was measured using the Enspire 2300 Multimode Plate Reader (PerkinElmer; Waltham, MA, USA) at an excitation wavelength of 530 nm and an emission wavelength of 590 nm. The protein content of cell lysates was measured as described above. Intracellular cholesterol level was normalized to total protein concentration.

### Cellular uptake of poly(I:C)

To measure cellular uptake of poly(I:C), J774.1/JA-4 cells were treated with 1 μg/mL of poly(I:C) (HMW)-Rhodamine for 2 h at 37 °C. The cells were washed three times with PBS and stained with 1 μg/mL of Hoechst 33342. Fluorescence of rhodamine was detected at an excitation wavelength of 540 nm and an emission wavelength of 580 nm using the EnSpire 2300 Multimode Plate Reader. To observe the cellular uptake of poly(I:C), the cells were treated with 1 μg/mL of poly(I:C) (HMW)-Rhodamine for 2 h at 37 °C in 8-well glass chamber slides (Corning, Corning, NY, USA). Then, they were washed with PBS, and fixed with 4% (w/v) paraformaldehyde in PBS on ice for 15 min. Coverslips were mounted onto chamber slides using ProLong Gold Antifade Mountant with DAPI. Images were obtained by laser scanning confocal microscopy (LSM700, Carl ZEISS, Oberkochen, Germany).

### Small interfering RNA knockdown of TLR3

Stealth RNAi small interfering RNA (siRNA) targeting the TLR3 gene (5’-CCUGAUGAUCUUCCCUCUAACAUAA-3’) and Stealth RNAi siRNA negative control med GC Duplex #2 were obtained from Thermo Fisher Scientific. J774.1/JA-4 cells were seeded at 5 x 10^5^ cells/well in 12-well plates and transfected with each siRNA (20 nM) using Lipofectamine 3000 (Thermo Fisher Scientific) according to the manufacturer’s instructions. After a 24-h transfection, the cells were harvested and subjected to qPCR analysis.

### Statistical analysis

All quantitative data are shown as means ± standard deviation (SD). Statistical significance was determined using Student’s *t*-test. Comparisons between more than two groups were performed using one-way ANOVA with Tukey’s *post hoc* test. *p* < 0.05 was considered statistically significant.

### Data availability statement

All data are contained within this manuscript and supporting information. Requests for additional information can be directed to the corresponding author: Dr. Ko Fujimori

## Acknowledgments

We thank Dr. Fumio Amano (Osaka University of Pharmaceutical Sciences) for his valuable comments.

## Author contributions

A.K. and K.F. contributed to the experimental design and data analysis. A.K. and K.T. performed the experiments. A.K. and K.F. wrote the manuscript with agreement of K.T.

## Funding and additional information

This study was supported in part by JSPS KAKENHI (JP17K15529 to A.K.).

## Conflict of interest statement

The authors declare that they have no conflicts of interest with the contents of this article.

## Footnotes

dsRNA, double-stranded RNA; TLR, Toll-like receptor; RLR, RIG-I-like receptor; RIG-I, retinoic acid-inducible protein-I; MDA5, melanoma differentiation-associated gene 5; poly(I:C), polyinosinic-polycytidylic acid; IFN, interferon; ISG, interferon-stimulated gene; HMG-CoA, 3-hydroxy-3-methyl glutaryl coenzyme A; LFD, low-fat diet; HFD, high-fat diet; NEFA, non-esterified fatty acids; LDL, low-density lipoprotein; HDL, high-density lipoprotein; IRF interferon regulatory factor; NF-κB, nuclear factor-κB; IL, interleukin; TNF, tumor necrosis factor; CCL, chemokine (C-C motif) ligand; RANTES, regulated on activation, normal T cell expressed and secreted; JAK, Janus kinase; STAT, signal transducers and activators of transcription; ISGF, interferon-stimulated gene factor; GGPP, geranylgeranyl pyrophosphate; GAPDH, glyceraldehyde-3-phosphate dehydrogenase; HRP, horseradish peroxidase; FBS, fetal bovine serum; siRNA; small interfering RNA; ROCK, Rho-associated, coiled-coil-containing protein kinase

## References

1. Endo, A., Kuroda, M., and Tsujita, Y. (1976) ML-236A, ML-236B, and ML-236C, new inhibitors of cholesterogenesis produced by Penicillium citrinium. J. Antibiot. (Tokyo) 29, 1346–1348

2. Goldstein, J. L., and Brown, M. S. (1990) Regulation of the mevalonate pathway. Nature 343, 425–430

3. Grundy, S. M., Stone, N. J., Bailey, A. L., Beam, C., Birtcher, K. K., Blumenthal, R. S., Braun, L. T., de Ferranti, S., Faiella-Tommasino, J., Forman, D. E., Goldberg, R., Heidenreich, P. A., Hlatky, M. A., Jones, D. W., Lloyd-Jones, D., Lopez-Pajares, N., Ndumele, C. E., Orringer, C. E., Peralta, C. A., Saseen, J. J., Smith, S. C., Jr., Sperling, L., Virani, S. S., and Yeboah, J. (2019) 2018 AHA/ACC/AACVPR/AAPA/ABC/ACPM/ADA/AGS/APhA/ASPC/NLA/PCNA guideline on the management of blood cholesterol: A report of the American college of cardiology/American heart association task force on clinical practice guidelines. J. Am. Coll. Cardiol. 73, e285–e350

4. Oesterle, A., Laufs, U., and Liao, J. K. (2017) Pleiotropic effects of statins on the cardiovascular system. Circ. Res. 120, 229–243

5. Ortego, M., Bustos, C., Hernández-Presa, M. A., Tuñón, J., Díaz, C., Hernández, G., and Egido, J. (1999) Atorvastatin reduces NF-κB activation and chemokine expression in vascular smooth muscle cells and mononuclear cells. Atherosclerosis 147, 253–261

6. Arnaud, C., Burger, F., Steffens, S., Veillard, N. R., Nguyen, T. H., Trono, D., and Mach, F. (2005) Statins reduce interleukin-6-induced C-reactive protein in human hepatocytes: new evidence for direct antiinflammatory effects of statins. Arterioscler. Thromb. Vasc. Biol. 25, 1231–1236

7. Tuomisto, T. T., Lumivuori, H., Kansanen, E., Häkkinen, S. K., Turunen, M. P., van Thienen, J. V., Horrevoets, A. J., Levonen, A. L., and Ylä-Herttuala, S. (2008) Simvastatin has an anti-inflammatory effect on macrophages via upregulation of an atheroprotective transcription factor, Kruppel-like factor 2. Cardiovasc. Res. 78, 175–184

8. Lee, C. S., Yi, E. H., Lee, J. K., Won, C., Lee, Y. J., Shin, M. K., Yang, Y. M., Chung, M. H., Lee, J. W., Sung, S. H., and Ye, S. K. (2013) Simvastatin suppresses RANTES-mediated neutrophilia in polyinosinic-polycytidylic acid-induced pneumonia. Eur. Respir. J. 41, 1147–1156

9. Akira, S., Uematsu, S., and Takeuchi, O. (2006) Pathogen recognition and innate immunity. Cell 124, 783–801

10. Thompson, A. J., and Locarnini, S. A. (2007) Toll-like receptors, RIG-I-like RNA helicases and the antiviral innate immune response. Immunol. Cell Biol. 85, 435–445

11. Meylan, E., and Tschopp, J. (2006) Toll-like receptors and RNA helicases: two parallel ways to trigger antiviral responses. Mol. Cell 22, 561–569

12. Wong, M. T., and Chen, S. S. (2016) Emerging roles of interferon-stimulated genes in the innate immune response to hepatitis C virus infection. Cell. Mol. Immunol. 13, 11–35

13. Mesev, E. V., LeDesma, R. A., and Ploss, A. (2019) Decoding type I and III interferon signalling during viral infection. Nat. Microbiol. 4, 914–924

14. Kraus, T. A., Lau, J. F., Parisien, J. P., and Horvath, C. M. (2003) A hybrid IRF9-STAT2 protein recapitulates interferon-stimulated gene expression and antiviral response. J. Biol. Chem. 278, 13033–13038

15. Schindler, C., Levy, D. E., and Decker, T. (2007) JAK-STAT signaling: from interferons to cytokines. J. Biol. Chem. 282, 20059–20063

16. Lazear, H. M., Pinto, A. K., Vogt, M. R., Gale, M., Jr., and Diamond, M. S. (2011) Beta interferon controls West Nile virus infection and pathogenesis in mice. J. Virol. 85, 7186–7194

17. Luker, G. D., Prior, J. L., Song, J., Pica, C. M., and Leib, D. A. (2003) Bioluminescence imaging reveals systemic dissemination of herpes simplex virus type 1 in the absence of interferon receptors. J. Virol. 77, 11082–11093

18. Hofer, M. J., Li, W., Manders, P., Terry, R., Lim, S. L., King, N. J., and Campbell, I. L. (2012) Mice deficient in STAT1 but not STAT2 or IRF9 develop a lethal CD4^+^ T-cell-mediated disease following infection with lymphocytic choriomeningitis virus. J. Virol. 86, 6932–6946

19. McKay, A., Leung, B. P., McInnes, I. B., Thomson, N. C., and Liew, F. Y. (2004) A novel anti-inflammatory role of simvastatin in a murine model of allergic asthma. J. Immunol. 172, 2903–2908

20. Sparrow, C. P., Burton, C. A., Hernandez, M., Mundt, S., Hassing, H., Patel, S., Rosa, R., Hermanowski-Vosatka, A., Wang, P. R., Zhang, D., Peterson, L., Detmers, P. A., Chao, Y. S., and Wright, S. D. (2001) Simvastatin has anti-inflammatory and antiatherosclerotic activities independent of plasma cholesterol lowering. Thromb. Vasc. Biol. 21, 115–121

21. Niessner, A., Steiner, S., Speidl, W. S., Pleiner, J., Seidinger, D., Maurer, G., Goronzy, J. J., Weyand, C. M., Kopp, C. W., Huber, K., Wolzt, M., and Wojta, J. (2006) Simvastatin suppresses endotoxin-induced upregulation of toll-like receptors 4 and 2 in vivo. Atherosclerosis 189, 408–413

22. Koike, A., Hanatani, M., and Fujimori, K. (2019) Pan-caspase inhibitors induce necroptosis via ROS-mediated activation of mixed lineage kinase domain-like protein and p38 in classically activated macrophages. Exp. Cell Res. 380, 171–179

23. Lacy, M., Atzler, D., Liu, R., de Winther, M., Weber, C., and Lutgens, E. (2019) Interactions between dyslipidemia and the immune system and their relevance as putative therapeutic targets in atherosclerosis. Pharmacol. Ther. 193, 50–62

24. Fatkhullina, A. R., Peshkova, I. O., and Koltsova, E. K. (2016) The role of cytokines in the development of atherosclerosis. Biochemistry Mosc. 81, 1358–1370

25. Slater, L., Bartlett, N. W., Haas, J. J., Zhu, J., Message, S. D., Walton, R. P., Sykes, A., Dahdaleh, S., Clarke, D. L., Belvisi, M. G., Kon, O. M., Fujita, T., Jeffery, P. K., Johnston, S. L., and Edwards, M. R. (2010) Co-ordinated role of TLR3, RIG-I and MDA5 in the innate response to rhinovirus in bronchial epithelium. PLoS Pathog. 6, e1001178

26. Mogensen, T. H. (2018) IRF and STAT transcription factors - From basic biology to roles in infection, protective immunity, and primary immunodeficiencies. Front. Immunol. 9, 3047

27. Black, S., Nicolay, U., Del Giudice, G., and Rappuoli, R. (2016) Influence of statins on influenza vaccine response in elderly individuals. J. Infect. Dis. 213, 1224–1228

28. McLean, H. Q., Chow, B. D., VanWormer, J. J., King, J. P., and Belongia, E. A. (2016) Effect of statin use on influenza vaccine effectiveness. J. Infect. Dis. 214, 1150–1158

29. Omer, S. B., Phadke, V. K., Bednarczyk, R. A., Chamberlain, A. T., Brosseau, J. L., and Orenstein, W. A. (2016) Impact of statins on influenza vaccine effectiveness against medically attended acute respiratory illness. J. Infect. Dis. 213, 1216–1223

30. Kim, M. C., Yun, S. C., Lee, S. O., Choi, S. H., Kim, Y. S., Woo, J. H., and Kim, S. H. (2018) Statins increase the risk of herpes zoster: A propensity score-matched analysis. PloS One 13, e0198263

31. Fan, L., Wang, Y., Liu, X., and Guan, X. (2019) Association between statin use and herpes zoster: systematic review and meta-analysis. BMJ open 9, e022897

32. Kwak, B., Mulhaupt, F., Veillard, N., Pelli, G., and Mach, F. (2001) The HMG-CoA reductase inhibitor simvastatin inhibits IFN-γ induced MHC class II expression in human vascular endothelial cells. Swiss Med. Wkly. 131, 41–46

33. Cole, J. E., Navin, T. J., Cross, A. J., Goddard, M. E., Alexopoulou, L., Mitra, A. T., Davies, A. H., Flavell, R. A., Feldmann, M., and Monaco, C. (2011) Unexpected protective role for Toll-like receptor 3 in the arterial wall. Proc. Natl. Acad. Sci. U.S.A. 108, 2372–2377

34. Tate, M. D., Pickett, D. L., van Rooijen, N., Brooks, A. G., and Reading, P. C. (2010) Critical role of airway macrophages in modulating disease severity during influenza virus infection of mice. J. Virol. 84, 7569–7580

35. Wheeler, D. L., Sariol, A., Meyerholz, D. K., and Perlman, S. (2018) Microglia are required for protection against lethal coronavirus encephalitis in mice. J. Clin. Invest. 128, 931–943

36. Okazaki, T., Higuchi, M., Takeda, K., Iwatsuki-Horimoto, K., Kiso, M., Miyagishi, M., Yanai, H., Kato, A., Yoneyama, M., Fujita, T., Taniguchi, T., Kawaoka, Y., Ichijo, H., and Gotoh, Y. (2015) The ASK family kinases differentially mediate induction of type I interferon and apoptosis during the antiviral response. Sci. Signal. 8, ra78

37. Calvi, S. A., Soares, A. M., Peraçoli, M. T., Franco, M., Ruiz, R. L., Jr., Marcondes-Machado, J., Fecchio, D., Mattos, M. C., and Mendes, R. P. (2003) Study of bronchoalveolar lavage fluid in paracoccidioidomycosis: cytopathology and alveolar macrophage function in response to gamma interferon; comparison with blood monocytes. Microbes Infect. 5, 1373–1379

38. Meyer, K. C., Raghu, G., Baughman, R. P., Brown, K. K., Costabel, U., du Bois, R. M., Drent, M., Haslam, P. L., Kim, D. S., Nagai, S., Rottoli, P., Saltini, C., Selman, M., Strange, C., and Wood, B. (2012) An official American thoracic society clinical practice guideline: the clinical utility of bronchoalveolar lavage cellular analysis in interstitial lung disease. Am. J. Respir. Crit. Care Med. 185, 1004–1014

39. Ourthiague, D. R., Birnbaum, H., Ortenlöf, N., Vargas, J. D., Wollman, R., and Hoffmann, A. (2015) Limited specificity of IRF3 and ISGF3 in the transcriptional innate-immune response to double-stranded RNA. J. Leukoc. Biol. 98, 119–128

40. Brandelius, A., Mahmutovic Persson, I., Calvén, J., Bjermer, L., Persson, C. G., Andersson, M., and Uller, L. (2013) Selective inhibition by simvastatin of IRF3 phosphorylation and TSLP production in dsRNA-challenged bronchial epithelial cells from COPD donors. Br. J. Pharmacol. 168, 363–374

41. Blanco-Colio, L. M., Tuñón, J., Martín-Ventura, J. L., and Egido, J. (2003) Antiinflammatory and immunomodulatory effects of statins. Kidney Int. 63, 12–23

42. Zhang, F. L., and Casey, P. J. (1996) Protein prenylation: molecular mechanisms and functional consequences. Annu. Rev. Biochem. 65, 241–269

43. Terao, Y., Satomi-Kobayashi, S., Hirata, K., and Rikitake, Y. (2015) Involvement of Rho-associated protein kinase (ROCK) and bone morphogenetic protein-binding endothelial cell precursor-derived regulator (BMPER) in high glucose-increased alkaline phosphatase expression and activity in human coronary artery smooth muscle cells. Cardiovasc. Diabetol. 14, 104

